# The ventral hippocampus and nucleus accumbens as neural substrates for cocaine contextual memory reconsolidation

**DOI:** 10.1101/2023.11.29.569314

**Authors:** Carolina Caban Rivera, Rachael Price, Ricardo P. Fortuna, Chen Li, Chau Do, Justin Shinkle, Marco G. Ghilotti, Xiangdang Shi, Lynn G. Kirby, George M. Smith, Ellen M. Unterwald

## Abstract

Drug craving triggered by cues that were once associated with drug intoxication is a major contributor to continued drug-seeking behaviors. Addictive drugs engage molecular pathways of associative learning and memory. Reactivated memories are vulnerable to disruption by interference with the process of reconsolidation, hence targeting reconsolidation could be a strategy to reduce cue-induced drug craving and relapse. Here we examined the circuitry of cocaine contextual memory reconsolidation and explored neuroplasticity following memory reactivation. Mice underwent chemogenetic inhibition of either nucleus accumbens (NA) neurons or the glutamatergic projection neurons from the ventral hippocampus (vHPC) to NA using inhibitory designer receptors exclusively activated by designer drugs (iDREADD). Mice underwent cocaine conditioned place preference followed by reactivation of the cocaine contextual memory. Clozapine-N-oxide (CNO) was administered after memory reactivation to inhibit either NA neurons or the accumbens–projecting vHPC neurons during the reconsolidation period. When retested 3 days later, a significant reduction in the previously established preference for the cocaine context was found in both conditions. FosTRAP2-Ai14 mice were used to identify neurons activated by cocaine memory recall and to evaluate plasticity in NA medium spiny neurons (MSNs) and vHPC pyramidal neurons upon recall of cocaine memories. Results indicate a significant increase in dendritic spine density in NA MSNs activated by cocaine memory recall, particularly of the thin spine type. Sholl analysis indicated longer dendritic length and more branching of NA MSNs after cocaine memory recall than without memory reactivation. vHPC neurons showed increased spine density, with the most robust change in stubby spines. These results implicate a circuit involving glutamatergic projections from the vHPC onto NA neurons which is necessary for the reconsolidation of cocaine memories. Interruption of cocaine memory reconsolidation reduced drug-seeking behavior.

## INTRODUCTION

Addictive drugs, including cocaine, engage molecular signaling pathways that are involved in associative learning processes. Cocaine-associated memories are highly resistant to extinction and exposure to cues previously associated with cocaine availability can elicit a conditioned physiological response, intense drug craving, and relapse to cocaine use [1, 2]. Once drug reward memories are reactivated, they undergo a process of reconsolidation through synaptic plasticity which strengthens the memory [3, 4]. During reactivation, memory traces are labile and can be manipulated behaviorally or pharmacologically [5, 6]. Since drug-associated cues can trigger craving and relapse, pharmacological disruption of reconsolidation-related plasticity that serves to maintain intrusive cocaine-related memories may be a useful approach to prevent relapse [7]. Indeed, interference with reconsolidation after cue re-exposure can reduce craving for cocaine and other drugs, at least transiently [7, 8]. Since exposure to drug cues can trigger relapse, a treatment objective is to extinguish previously learned associations between the positive effects of cocaine and environmental cues signaling cocaine availability [7, 9–13]. The development of successful therapeutic strategies for dampening cue-induced craving is dependent on a greater understanding of the circuitry and molecular processes involved in maintenance of cocaine memories.

The goal of this study was to identify the neural circuitry engaged in cocaine memory maintenance and elucidate specific cell populations required for reconsolidation of cocaine contextual memories. The literature supports the involvement of the ventral hippocampus (vHPC) in circuits of fear-memory reconsolidation [14–16]. There also is support for the involvement of projections from the vHPC to the nucleus accumbens (NA) in the retrieval of recent cocaine memory [17]. Other work indicates the NA as an important site in the maintenance of drugs memories [18–20]. Our previous work identified a signaling pathway consisting of NMDA receptors→GSK3β→mTORC1 in the HPC and NA whose activity is required for reconsolidation of cocaine contextual memories [21–23]. Given these findings, we hypothesized that the maintenance of cocaine reward memories relies on activation of neurons in the NA as well as their afferent connections from the vHPC. As such, we anticipated that inhibition of the vHPC→NA circuit after cocaine memory recall would abolish a previously established preference for a cocaine-paired environment. To test this, we utilized mice expressing pathway-targeted iDREADDs to silence specific neurons specifically during reconsolidation of cocaine reward memory. We characterized cell types activated in the NA and vHPC by the recall of cocaine contextual memories and examined dendritic spine plasticity in these active neurons.

## MATERIALS AND METHODS

### Animals

Animal procedures were performed in compliance with the National Institutes of Health guidelines for the Care and Use of Laboratory Animals, and animal use was reviewed and approved by Temple University Institutional Animal Care and Use Committee. Male C57BL/6 (8 weeks old on delivery, Charles River Laboratories, Wilmington, MA) were single housed under a 12-hour light/dark cycle (7:00 AM/7:00 PM) without enrichment objects with ad libitum standard chow and water. Animals were housed for five days before experiments began and weighed daily. Behavioral procedures were conducted between 9:00-11:00. Some studies utilized male and female FosTRAP2-Ai14 mice, 8-12 weeks old. FosTRAP2 mice were crossed with Ai14 reporter mice (expressing TdTomato (TdT)), both purchased from Jackson Labs (Stock 030323 & 007914) and bred in our facility. TRAP2 mice utilize a Fos promoter to drive transcription of CreER permitting temporally controlled recombination in active neurons when 4-hydroxytamoxifen (4-OHT) is administered. With the TRAP2-Ai14 cross, 4-OHT leads to the permanent expression of TdT in Fos-expressing neurons.

### Drugs

Cocaine hydrochloride, generously supplied by the National Institute on Drug Abuse Drug Supply Program, and clozapine-N-oxide hydrochloride (CNO; Cayman Chemicals, #25780) were dissolved in sterile saline. 4-Hydroxytamoxifen (4-OHT; Sigma #H6278) was dissolved in corn oil according to a modification of published methods [24]. All drugs were injected intraperitoneally (ip) in a volume of 5 ml/kg body weight and equal volumes of saline or corn oil served as vehicle controls.

### Viral Vectors

A chemogenetic approach allowed for selective expression of iDREADDs and neuronal inhibition. In experiment 1, iDREADDs were expressed in the nucleus accumbens (NA) neuronal cell bodies using AAV2-hSyn-hm4Di-mCherry. In experiment 3, glutamatergic neuronal cell in the ventral hippocampus (vHPC) projecting to the NA were targeted using a two-vector system approach. In this case, AAV5-calcium/calmodulin dependent protein kinase II-cre was delivered into the bilateral vHPC, and a Cre-dependent retrograde iDREADD (retroAAV2-DIO-hM4D(Gi)-mCherry) was delivered into the bilateral NA. The AAV vectors were obtained from the Neural Repair Viral Core (Shriners grant #84051-PHI-21, GM Smith), where they were fully characterized and tested for specificity.

Recombinant adeno-associated viruses were generated using pAAV-hSyn-hM4D(Gi)-mCherry or pAAV-hSyn-DIO-hM4D(Gi)-mCherry (gifts from Brian Roth (Addgene #50475 and #44362)) and pAAV-CamKII-Cre (gift from Janelia Viral Tools (Addgene #182736)) by a helper virus-free system [25]. 293T cells were grown to 70-80% confluence at which point they were transfected with two packaging plasmids using PEI (Polyethylenimine, linear, MW-25k, Warrington, PA): one carrying the AAV rep, cap genes (either AAV2, AAV5 or AAVretro) and a helper plasmid carrying the adenovirus helper functions. Three days after transfection, the cell lysates and the supernatant were harvested and 40% PEG 8000 was added to precipitate crude virus for 2 hours. Then the viruses were purified by double-centrifugation with cesium chloride (CsCl) and the isolated virus was dialyzed in 0.1M PBS/5% sorbital overnight [25, 26]. The viral titer of AAV2-hSyn-hM4D(Gi)-mCherry (4.10x1011 GC/mL), AAVretro-hSyn-DIO-hM4D(Gi)-mCherry (1.02x1014 GC/mL), AAVretro-hSyn-mCherry (3.67x10^13 GC/mL) and AAV2-CamKII-cre (1.0x1012 GC/mL) were determined by quantitative real-time PCR and expression was tested by transduction of fibroblasts and imaging for red fluorescence or Cre activity.

### Intracranial Vector Injections

Surgical procedures were carried out to inject viral vectors into the NA and vHPC as described previously by us [22, 27]. On the day of stereotaxic surgery, mice were anesthetized with isoflurane and secured into a stereotaxic frame (Stoelting Co.) fitted with a gaseous anesthesia nosecone delivering isoflurane. Using sterile instruments and techniques, an incision was made, and the skin was retracted over the top of the skull. A small hole was drilled through the skull over the site of interest, which was targeted bilaterally. Using the stereotaxic apparatus, an injector attached to a Hamilton syringe and syringe pump was lowered into the brain using the following coordinates [27]: NA (AP: +0.85 mm; ML: ±0.70 mm; DV: -4.2 mm); vHPC (AP: - 3.3, ML: ±3.0; DV: -3.0 and -4.0). The incision was closed with wound clips or surgical glue, and the analgesic agent meloxicam was administered post-surgery. Mice were monitored by the surgeon for recovery from anesthesia and general health before being single housed for recovery and experiment. Meloxicam administration was repeated the day after surgery. The functionality of the hM4Di synthetic receptor in the presence of CNO was verified ex vivo using whole-cell patch-clamp slice electrophysiology (data not shown).

### Cocaine-induced conditioned place preference (CPP)

An unbiased procedure was used following our published methods [21–23]. Place conditioning occurred in rectangular plastic chambers (45 x 20 x 20 cm) consisting of two unique compartments, one with white and black vertical striped walls and smooth flooring and the other with white walls with black circles and rough flooring. The two compartments were separated by a removable wall during conditioning and a door during testing. Illumination in both compartments was equal and mice showed no initial preference to either compartment. An unbiased procedure was used. Mice were injected with 10 mg/kg ip of cocaine or saline and were immediately confined to one compartment of the conditioning chamber, where they remained for 30 minutes. Conditioning occurred once per day for 8 consecutive days, resulting in four conditioning sessions with saline in one compartment, and four sessions with cocaine in the opposite compartment of the conditioning chamber, in a counterbalanced design. The test for place preference occurred on day 9, when mice had access to both compartments for 15 minutes in a drug-free state. The time spent in each compartment was recorded. Preference scores were calculated as: (time spent in the cocaine-paired compartment) minus (time spent in the saline-paired compartment) and reported in seconds. Mice that did not show a cocaine place preference on day 9 did not continue in the study. On day 10, 24 hours following assessment of cocaine place preference, some mice were re-exposed to the previously cocaine-paired compartment for 10 minutes to reactivate cocaine-associated memory, while other mice remained in the home cage.

### Chemogenetic neuronal inhibition

Mice expressing iDREADDs underwent cocaine CPP. On day 10, mice received CNO (5 mg/kg ip) or saline immediately after the 10 minute re-exposure to the cocaine context, and again 2 and 4 h post re-exposure to inhibit neuronal activity for the entire duration of reconsolidation (∼4-6 h) [28–30]. Mice were re-tested for cocaine place preference in a drug-free state 3 days post re-exposure. As a control for memory re-activation, some mice received an injection of CNO, same dose and schedule, without memory reactivation; they remained in home cage on day 10 and received CNO or vehicle injection in home cage.

### Immunohistochemistry

Following behavioral testing, mice underwent transcardiac perfusions and brain extractions. The brains were frozen and cryosectioned for immunostaining to verify iDREADD location, using primary antibody anti-RFP (1:1000, #600-401-379, Rockland). Sections were imaged at 10x using the Nikon AR2 confocal microscope.

### Dendritic Spine Analysis

Male and female FosTRAP2-Ai14 mice underwent an unbiased cocaine CPP procedure as described above. On day 10, mice were re-exposed to the cocaine compartment for 10 min to reactivate the cocaine memory and controls remained in their home cage. 4-OHT (50 mg/kg, i.p.) or vehicle (corn oil) was injected immediately after memory reactivation to “trap” cells that were activated by cocaine cue recall. After 7 days (maximal TdT expression), mice underwent transcardiac perfusions and brains were extracted. Brains were frozen and cryosectioned at 50 µm into free floating wells. IHC was conducted using rabbit primary antibody dsRed (1:500, #632496, Takara) to label TdT+ cells. Sections were imaged first at 10x using Nikon AR2 confocal microscope for localization of TdT labeled accumbens medium spiny neurons (MSN) and hippocampal pyramidal neurons, then individual neurons were imaged at 60x oil lens for detection of dendritic spines. After confocal imaging, cells were reconstructed using Neurolucida 360 software (Microbrightfield Bioscience). The cell soma was detected, then dendritic branches were traced using the directional kernels method. Spines were automatically detected on the reconstructed dendrites; corrections were manually made for accuracy. The tracing was exported to Neurolucida explorer. Cells were selected based on thorough expression of TdT, and dendrites free of obstructions from other cells or blood vessels that can be traced without interruption. Morphometrics were obtained using Neurolucida Explorer software. Parameters included total number of dendritic spines, head diameter, density per cell, morphology (mushroom, stubby, thin), and density per morphology, completed for 6-10 neurons per region per animal. Filipodia spines were not detected thus were not included in the analysis. Selected reconstructed cells from dendritic spine analysis were used to assess dendritic tree plasticity using Sholl analysis. Using the Neurolucida Explorer software, total dendritic length and number of dendritic intersections were counted in concentric circles of increasing diameter (10μm) from the nucleus of the cell.

## Results

### Inhibition of nucleus accumbens neuronal activity prevents memory reconsolidation and erases a previously established cocaine place preference

Experiment 1 determined if neurons in the nucleus accumbens are involved in cocaine contextual memory reconsolidation. Figure 1 summarizes the experimental design of the chemogenetic studies (Exp 1 and 3). AAV2-hSyn-hM4D(Gi)-mCherry was injected bilaterally into the accumbens prior to cocaine place conditioning and testing. Mice underwent 8 days of alternating cocaine and saline conditioning followed by a post-conditioning test for place preference on day 9. Figure 2A shows similar CPP scores on day 9 for the two groups of mice. On day 10, mice were re-exposed to the cocaine context for 10 min and CNO or vehicle administered immediately thereafter. Three days later, mice were re-tested for place preference. Preference scores from days 9 and 13 were analyzed by a two-way ANOVA with day and treatment factors (Fig 2A). Results indicate a signigicant interaction (F(1,12)=4.847; p=0.048) and main effect of day (F(1,12)=13.37; p=0.0033; treatment F(1,12) = 2.865; p=0.1163). Post hoc tests shows significantly lower CPP scores in mice expressing the iDREADD and injected with CNO following recall of the cocaine context, compared to mice injected with vehicle on Day 13 (iDREADD CNO vs vehicle) (p<0.05) and also significantly lower preference scores on day 13 vs day 9 for the iDREADD+CNO group (day 9 vs day 13, p<0.005). Figure 2B shows an example of bilateral expression of the iDREADD (AAV2-hSyn- hM4D(Gi)-mCherry) in the accumbens, corroborating site specific expression of iDREADDs. In a control experiment, inhibition of NA neurons by administration of CNO did not alter place preference when CNO was administered to mice in their home cage environment without re-exposure to the cocaine-paired context (Fig 2C). These data demonstrate that recall of cocaine memories was necessary for interferance of an established place preference. Together, these data suggest that inhibition of NA neurons after reactivation of a cocaine contextual memory, prevents reconsolidation as demonstrated by the loss in preference for the cocaine context 72 h post CNO administration.

**Figure 1.**
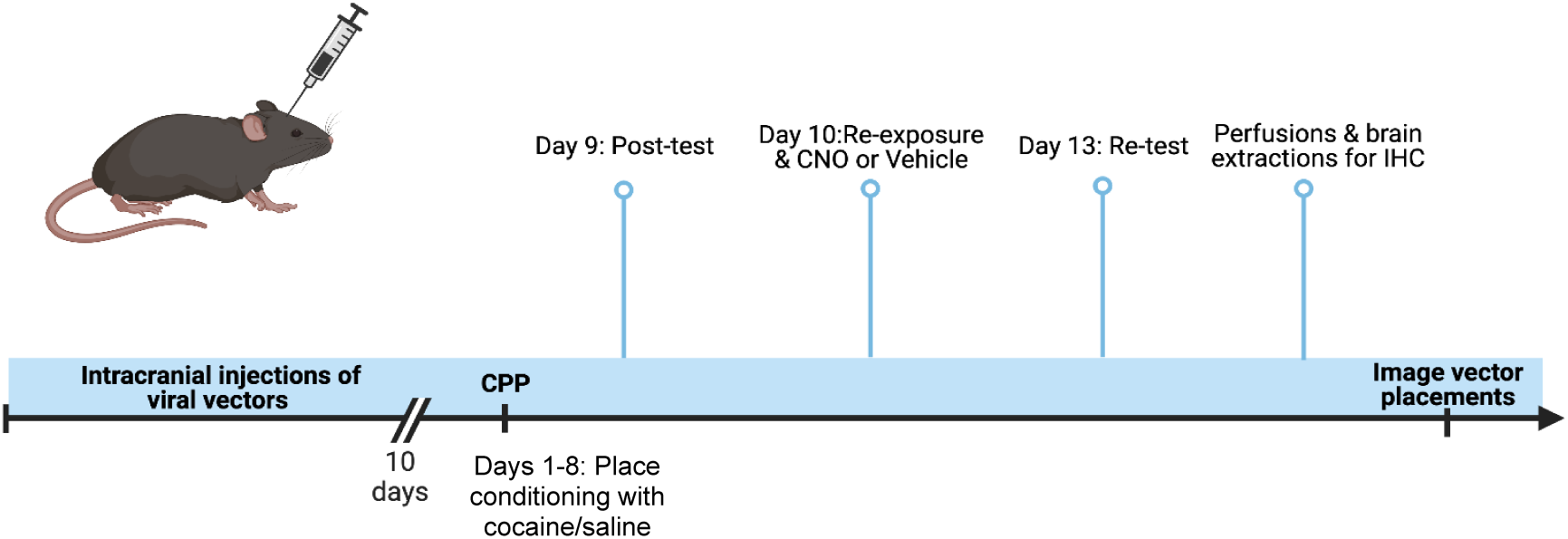
Experimental timeline of chemogenetic inhibition of targeted neurons during reconsolidation of cocaine contextual memory. For Experiments 1 and 3, mice received intracranial infusions of viral vectors 10-14 days prior to beginning cocaine CPP to allow stable expression of iDREADDs by the memory reactivation on Day 10 of CPP (ie, 20-24 days post viral injection). Created with BioRender.com.

**Figure 2.**
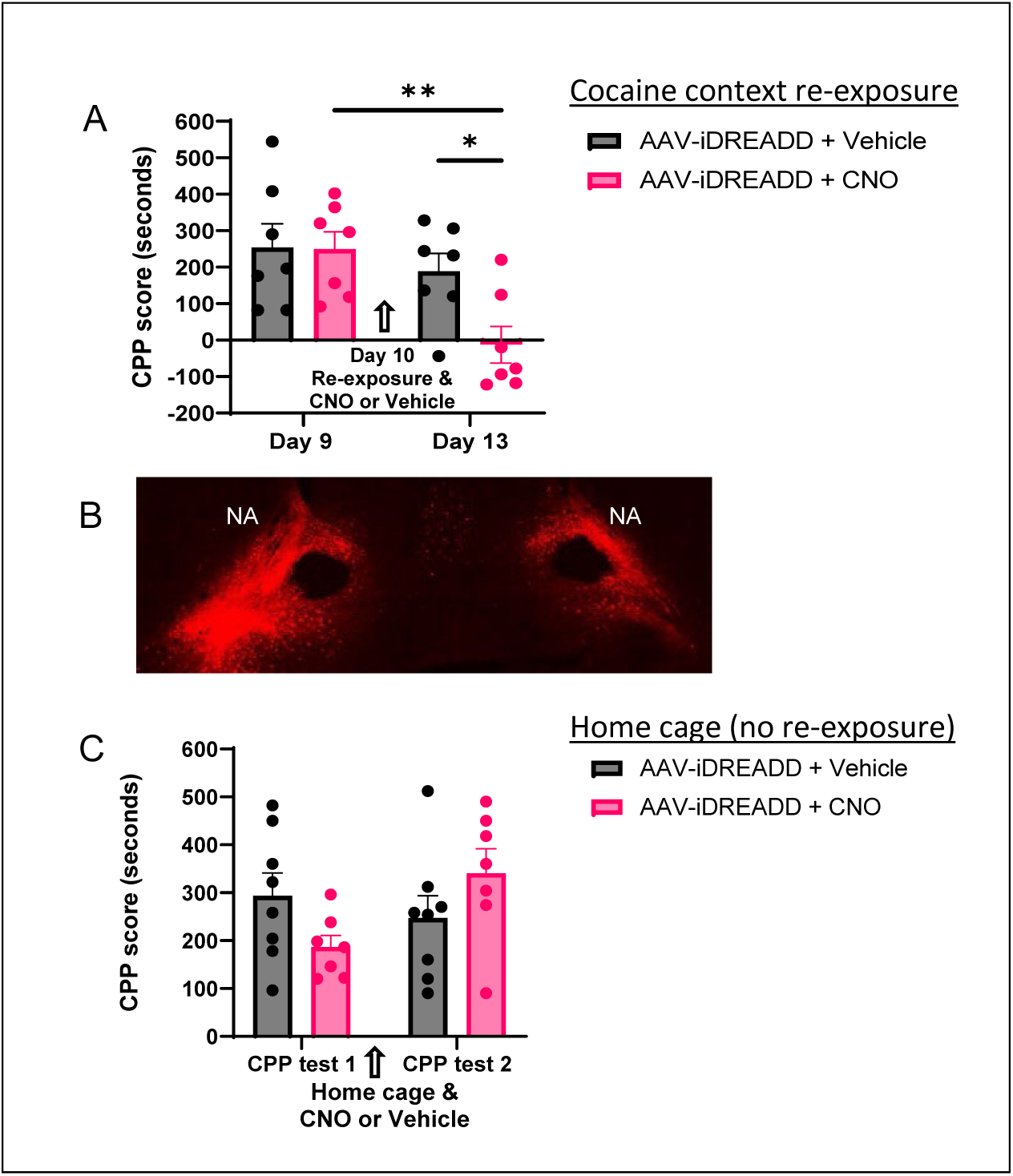
A) Inhibition of nucleus accumbens neurons during reconsolidation abolished cocaine place preference in subsequent testing. Two groups of mice expressing NA iDREADDs show similar cocaine (10 mg/kg) place preference on Day 9. On Day 10, mice were re-exposed to the cocaine context to activate memory and injected with vehicle or CNO (5 mg/kg) to inhibit NA neuronal firing. CPP was re-tested on Day 13. CPP scores (time in cocaine side – time in saline side, sec) were significantly lower in iDREADD-expressing mice injected with CNO compared to vehicle-injected mice on day 13 (*p<0.05) and significantly lower on day 13 than on day 9 (**p<0.01). (B) Representative images of nucleus accumbens (NA) showing expression of mCherry. (C) Two groups of mice expressing NA iDREADDs showed similar cocaine CPP scores on test 1. CNO (5 mg/kg) or vehicle was administered in the home cage, without cocaine context re-exposure, and preference retested 3 days later. No significant differences in CPP scores were found on test 2.

### Nucleus accumbens neurons are activated by recall of cocaine contextual memories

Experiment 2 utilized FosTRAP2-Ai14 mice to visualize and morphologically characterize neurons activated by recall of cocaine contextual memories. Two groups of mice underwent an unbiased cocaine CPP procedure. CPP scores are presented in Figure 3A; both groups had comparable preference for the cocaine-paired context on day 9. On day 10, one group of mice were re-exposed to the cocaine compartment (10 min, drug free) for reactivation of cocaine contextual memory and immediately thereafter administered 4-OHT (50 mg/kg) to label (ie “trap”) Fos+ cells with TdT. The other group of mice remained in their home cages and were injected with 4-OHT (50 mg/kg). Brains were prepared for analysis of TdT labeled cells in the nucleus accumbens. Figure 3B contains representative images of TdT+ cells in the accumbens after re-exposure to the cocaine context and for home cage controls. Both in the accumbens core and shell, reactivation of cocaine contextual memory led to an increase in the number of TdT+ cells compared to home cage controls (Figure 3C). Planned comparison of cell counts show a significant effect of re-exposure in the core (t=2.906, df=12, p=0.0132) and shell (t=4.205, df=12, p=0.0012) regions of the nucleus accumbens.

**Figure 3.**
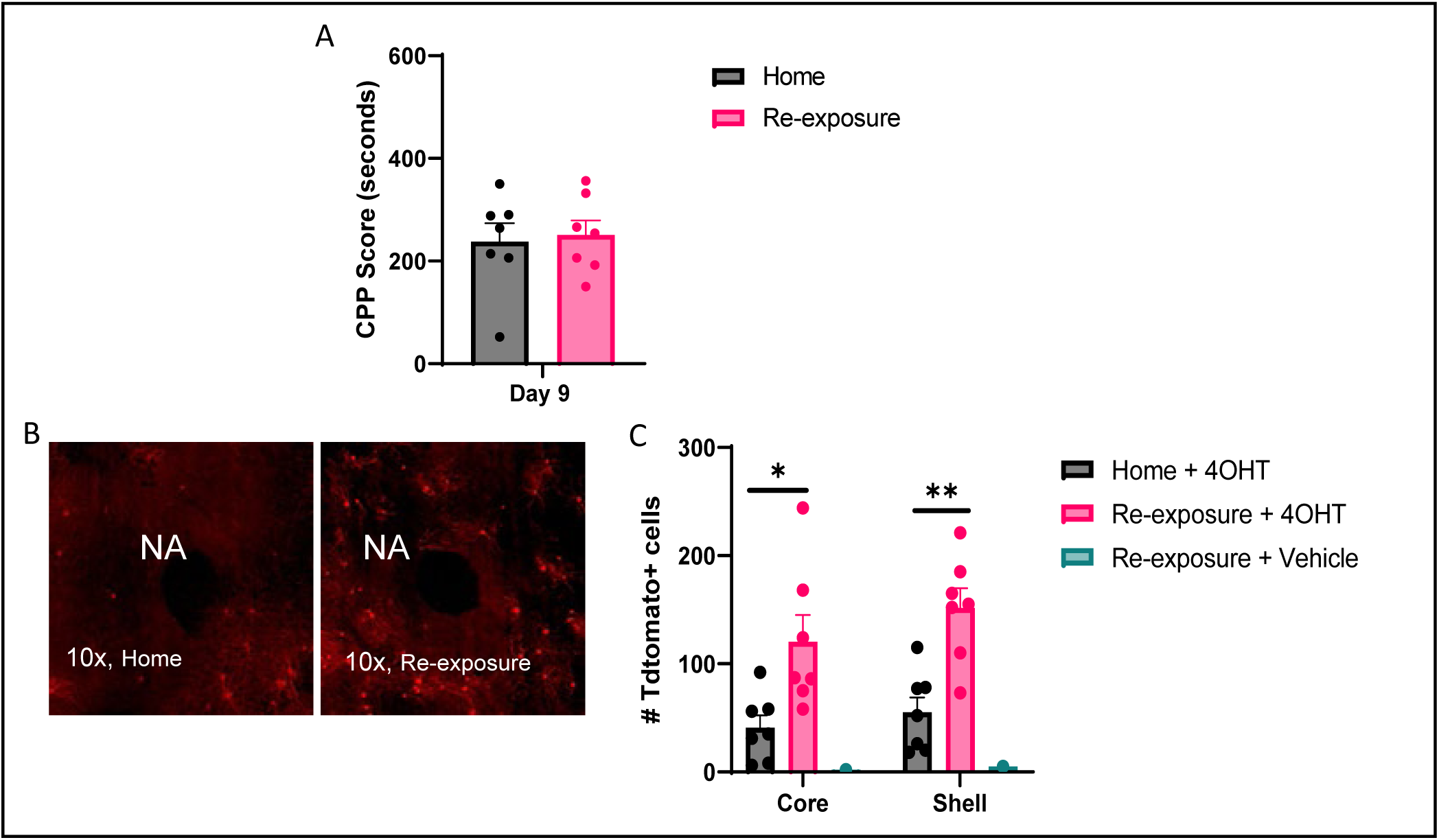
Quantification of TdT+ medium spiny neurons in the nucleus accumbens after cocaine context re-exposure. A) Cocaine place preference scores of two groups of TRAP2XAi14 mice showing both groups developed a preference for the cocaine-paired context. B) Representative images of TdT+ neurons following 4-OHT (50 mg/kg) administration in the home cage (left) or following re-exposure to the cocaine context (right). C) Quantification of TdT+ neurons showed a significantly greater number of active cells in the nucleus accumbens core (*p<0.05) and shell (**p<0.01) following recall of cocaine contextual memory compared to home cage controls.

### Dendritic spine plasticity in the accumbens following recall of cocaine contextual memories

Brains from the FosTRAP2-Ai14 mice used in Experiment 2 were further examined for synaptic plasticity following recall of cocaine memories. Medium spiny neurons (MSN) were identified by morphological examination of TdT+ cells and dendrites analyzed for spine density and characteristics. Figure 4A is a representative image of a TdT+ MSN at high magnification (left) along with a representative image of dendrite reconstruction and spine capture using Neurolucida 360 imaging software (right).

**Figure 4.**
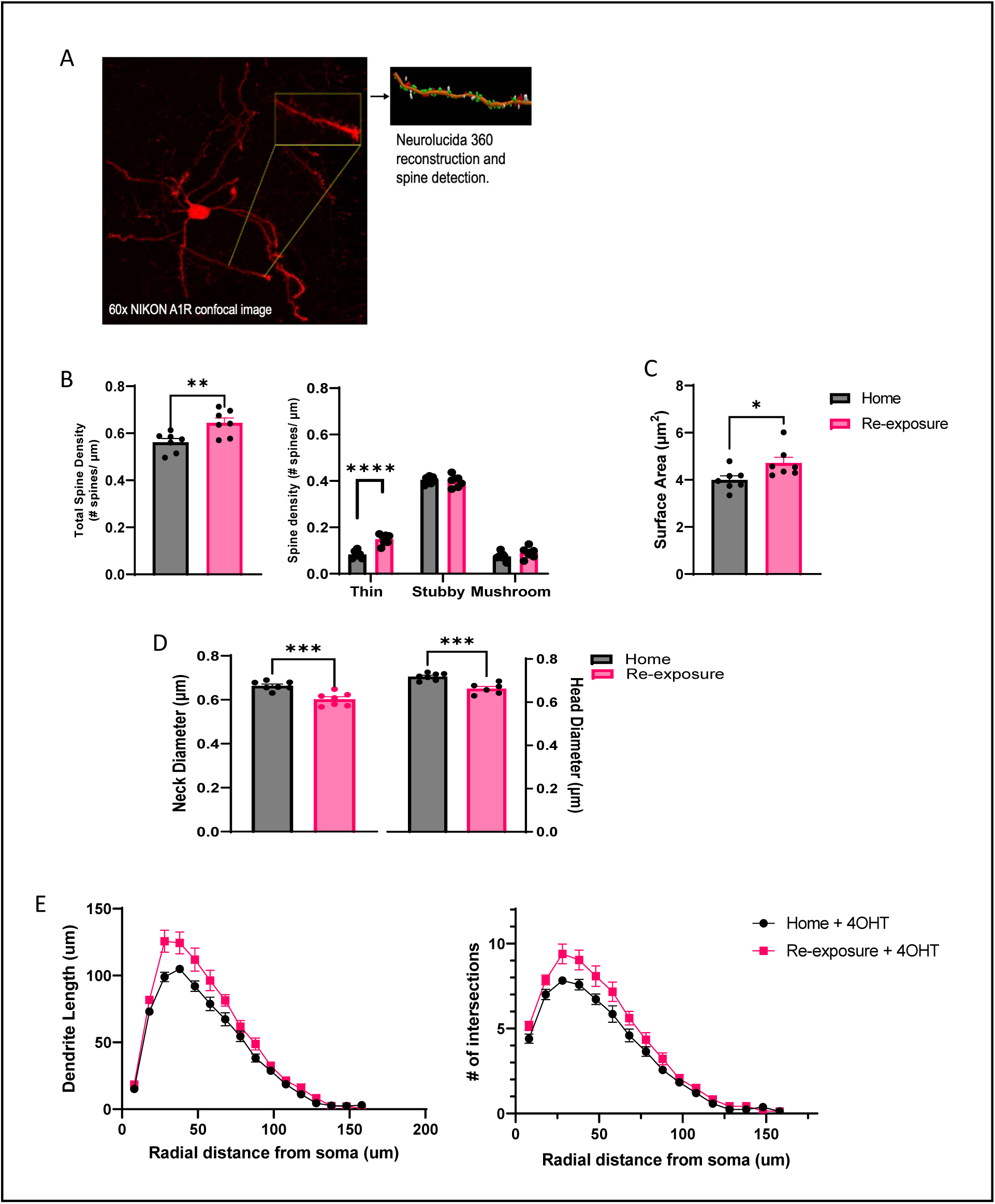
Dendritic spine analysis of TdT+ MSNs in the nucleus accumbens after cocaine context re-exposure. A) Representative 60x image of a TdT+ medium spiny neuron (left) and dendrite/spine capture from a reconstructed neuron (right). B) Quantification of total spine density (left) and spine density by morphology (right) following recall of cocaine contextual memory compared to home cage controls (**p<0.01, ****p<0.0001). Recall of cocaine contextual memory resulted in higher total spine density and higher density specifically of thin spines in accumbens medium spiny neurons. C) Quantification of spine surface area following recall of cocaine contextual memory compared to home cage controls (*p<0.05). D) Spine neck diameter (left) and head diameter (right) following recall of cocaine contextual memory were significantly smaller compared to home cage controls (***p<0.001). E) Sholl analysis of dendritic length (left) and number of intersections (right) showed that MSN dendrites were significantly longer (****p<0.0001) and more complex (****p<0.0001) following cocaine memory recall. 8-10 neurons/mouse were analyzed for morphometric characteristics; mean values/mouse are shown.

Results of the spine density analysis following re-exposure to cocaine context compared to home cage controls are shown in Figure 4B. Recall of cocaine contextual memory led to a significant increase in total spine density compared to home cage controls (t=3.137, df=12, p=0.0086) and this was driven by an increase in the density of thin spines (treatment factor F(1,35)=15.61, p=0.0004; morphology F(2,35)=10.33, p<0.0001; interaction F(2,35)=11.12, p=0.0002; post-hoc thin spines, p<0.0001). Total spine surface area was significantly higher following the recall of cocaine context as compared to no recall (Fig 4C; t=2.432, df=12; p=0.031). Spine neck diameter (left) and spine head diameter (right) were significantly smaller in dendrites of medium spiny neurons from mice re-exposed to the cocaine context as compared to home-cage controls (Fig. 4D; neck diameter t=4.549, df=12, p=0.0007; head diameter t=4.884, df=11, p=0.0005). Lower spine neck and head diameter is likely a reflection of the higher density of thin spines which have smaller head diameter than the other spine types. Sholl analysis indicated significant differences in dendritic length and complexity of MSNs of mice re-exposed to the cocaine context as compared with home cage controls (Fig 4E; two-way ANOVA length: treatment F(1,172)=19.59, p<0.0001; intersections: treatment F(1,162)=17.73, p<0.0001).

### Inhibition of glutamatergic projections from the ventral hippocampus to the nucleus accumbens prevents cocaine contextual memory reconsolidation

Cocaine memory reactivation caused significant neuroplasticity in the nucleus accumbens as demonstrated by changes in dendritic spines on the medium spiny neurons. Experiment 3 sought to identify the circuit that innervates the accumbens (NA) and contributes to the reconsolidation process. This study targeted the nucleus accumbens-projecting ventral hippocampal (vHPC) neurons and assessed their role in the reconsolidation of cocaine contextual memories. Figure 1 summarizes the experimental design. AAVretro-hSyn-DIO- hM4D(Gi) or control AAVretro-hSyn-mCherry vector was injected bilaterally into the nucleus accumbens and AAV-CamKII-Cre into the vHPC of mice, to express iDREADDs specifically in the vHPC→NA pathway. Mice underwent cocaine conditioning followed a test for place preference. Day 9 CPP scores are shown in Figure 5A. On day 10, cocaine contextual memory^N^w^A^as reactivated via a 10 min cocaine context re-exposure immediately followed by CNO or vehicle injection to activate the iDREADD. Results (Fig 5A) show that inhibition of vHPC→NA neurons by administration of CNO during reconsolidation abolished a previously established place preference when retested three days later on day 13. Mice expressing the control vector and injected with CNO maintained their cocaine place preference, as did mice expressing iDREADDs and injected with vehicle. CPP scores were analyzed by two-way ANOVA (treatment F(2,22)=5.440, p=0.012; day F(1,22)=31.44, p<0.0001; interaction F(2,22)=3.529, p=0.0468). Post-hoc analysis shows a significant difference of the rAAV- iDREADD+CNO group between day 9 and 13 (p=0.0001), and a significant difference between groups on day 13 (p=0.0111 rAAV-iDREADD+CNO vs rAAV-control+CNO; p=0.001 iDREADD+CNO vs iDREADD+vehicle). Thus the established cocaine preference was abolished by CNO-induced chemogenetic inhibition of the vHPC to NA glutamatergic projection. Location of vectors was verified and representative images of the NA injection site of the AAVretro-hSyn-DIO-hM4D(Gi)-mCherry and the vHPC expression of mCherry are provided in Figure 5B and 5C respectively.

**Figure 5.**
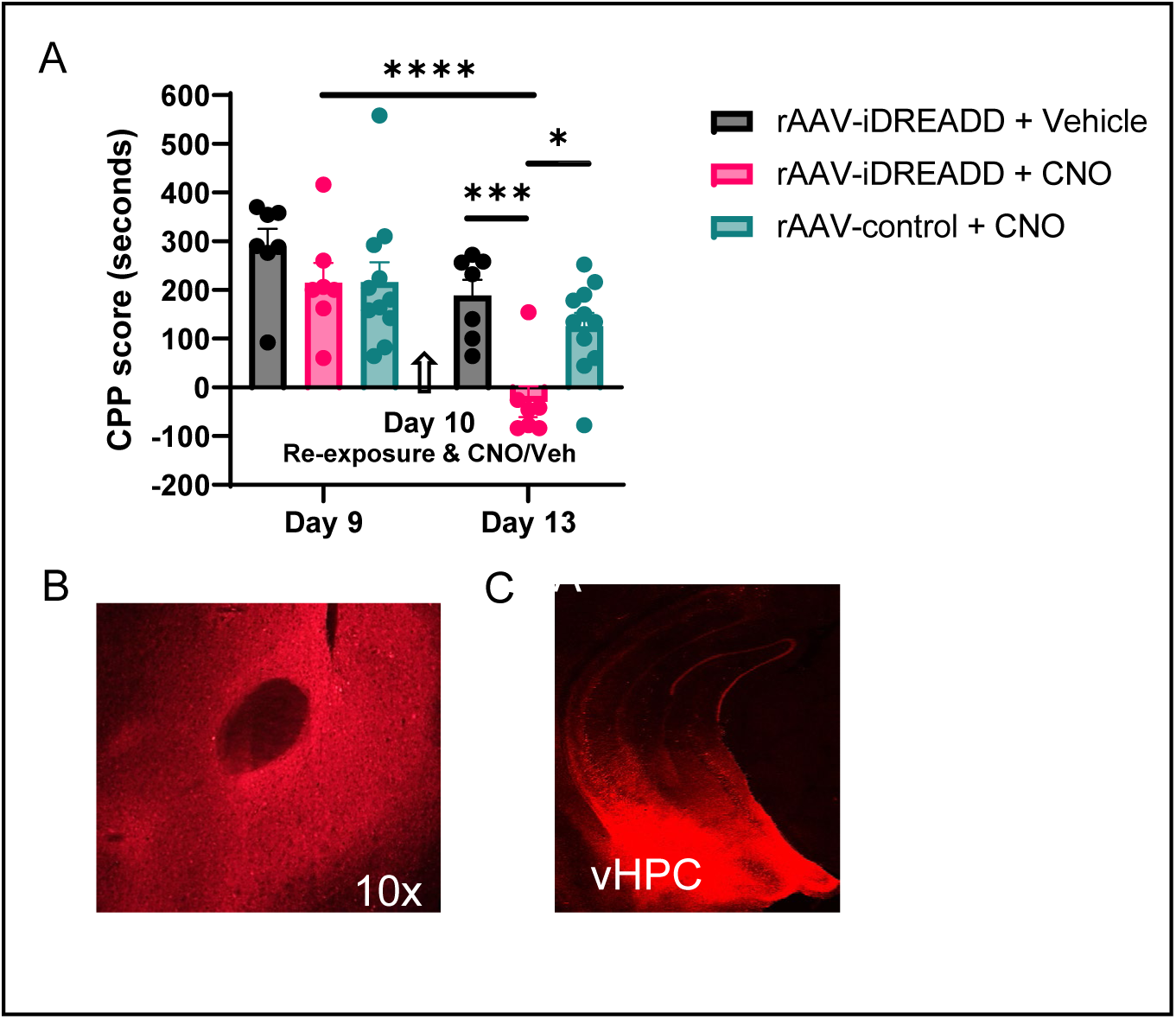
Inhibition of vHPC to NA glutamatergic projections disrupts reconsolidation of cocaine contextual memories. AAVretro-hSyn-DIO-hM4D(Gi) or control AAVretro-hSyn-mCherry vector was injected bilaterally into the nucleus accumbens and AAV-CamKII-Cre into the vHPC of mice, to express iDREADDs specifically in the vHPC→NA pathway. A) Mice expressing vHPC to NA iDREADDs or vector controls show similar cocaine CPP on Day 9. On day 10, mice were re-exposed to the cocaine context and injected with CNO (5 mg/kg) or vehicle. CPP was retested on day 13. CPP scores on day 13 were significantly lower in mice expressing iDREADDs and injected with CNO compared to both vehicle and vector control mice (*p<0.05; ***p<0.001). Day 13 scores were also significantly lower for the iDREADD + CNO group on day 13 versus day 9 (**** p<0.0001). B) Expression of mCherry in the NA. C) Expression of mCherry in the vHPC.

These data demonstrate that neurons in the vHPC were important in the maintenance of cocaine contextual memories. Neuroplasticity of these neurons was investigated similar to the analysis of the accumbens MSNs. Results demonstrate significant differences in dendritic spine density in vHPC pyramidal neurons following cocaine memory reactivation. Following establishment of a cocaine conditioned place preference, half the mice remained in their home cage while the others were re-exposed to the prior cocaine-paired context to reactivate cocaine contextual memories, followed by an injection of 4-OHT to ‘trap’ activated neurons. Analysis of TdTomato+ dendrites of ventral hippocampal pyramidal neurons demonstrated significantly greater spine density following recall of cocaine memories compared with home-cage controls as shown in Fig 6B (t= 4.903, df=9, p=0.0008). Analysis of spine density by morphology showed a significant effect of re-exposure (re-exposure F(1,27)=30.76, p<0.0001; morphology F(2,27)=1021, p<0.0001; interaction F(2,27)=0.5059, p=0.6086). Post-hoc test showed significant differences in the density of each spine type; thin spines (*p=0.0388), stubby spines (**p=0.0013), and mushroom spines (*p=0.0196) between home cage controls and cocaine context re-exposure. Total spine surface area in the vHPC was significantly higher following the recall of cocaine context as compared to no recall (Fig. 6C; t=9.373, df=9; p<0.0001). Spine neck diameter (Fig. 6D left; t=2.397 , df=9 , p=0.0401) and head diameter (Fig. 6D right; t=2.783, df=9, p=0.0213) were significantly larger in dendrites of pyramidal neurons from mice re-exposed to the cocaine context as compared to home cage controls. Spine volume was significantly greater in mice re-exposed to the cocaine context when compared to the home cage controls (Fig.6E; volume t=9.754, df=9, p<0.0001). Sholl analyses of dendritic length and complexity of pyramidal cells are shown in Figure 6F; no significant differences in dendritic length or intersections were found.

**Figure 6.**
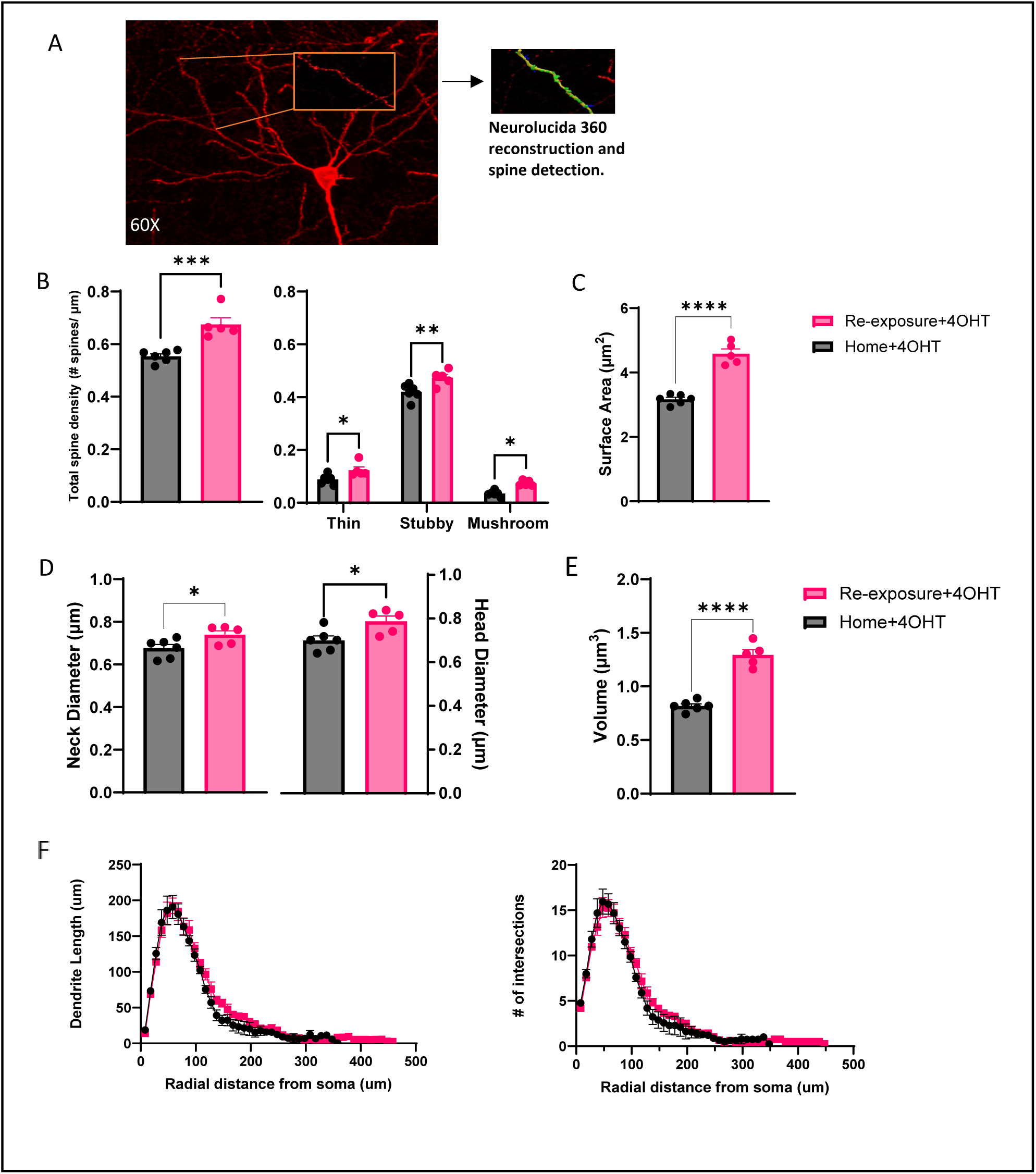
Dendritic spine analysis of TdT+ hippocampal pyramidal neurons after cocaine context re-exposure. A) Representative 60x image of a TdT+ pyramidal neuron in the vHPC (left) and dendrite/spine capture from a reconstructed neuron (right). B) Quantification of total spine density and spine density by morphology following recall of cocaine contextual memory compared to home cage controls. Recall of cocaine contextual memory resulted in higher total spine density (***p<0.001), due to greater density of all spine types (*p<0.05, **p<0.01). C) Spine surface area was significantly greater after cocaine memory recall as compared with home cage controls (****p<0.0001). D) Spine head diameter (*p<0.01) and neck diameter (*p<0.01) were larger after re-exposure to the cocaine context. E.) Spine volume (****<0.0001) was significantly greater in mice re-exposed to the cocaine context as compared to the home cage controls F) Sholl analysis of dendritic length and complexity are shown. 5- 8 neurons/mouse were analyzed for morphometric characteristics; mean values per mouse are shown.

## Discussion

Environmental cues can form strong and long-lasting associations with cocaine intoxication. This can occur when a context is paired with the rewarding effects of cocaine over multiple occasions. Repetition of such rewarding experience with a specific context leads to the context acquiring conditioned reinforcing properties [31–35]. Reactivation of drug memories that occurs when the contextual cues are present can trigger a physiological response and intense drug craving which can perpetuate drug-seeking behaviors. Disruption of cocaine contextual memories may be a valuable strategy to prevent cue-induced drug seeking behavior.

The present study sought to identify the circuit engram involved in the reconsolidation of cocaine memories. Our results demonstrate that cocaine memory reconsolidation can be disrupted by chemogenetically inhibiting NA intrinsic neurons or the glutamatergic projections from the vHPC to the NA. Inhibition of NA neuronal firing immediately after reactivation of cocaine-associated contextual memory erased a previously established cocaine place preference. NA neuronal silencing in the absence of memory recall (i.e., CNO to home cage controls) had no effect on the maintenance of the place preference, supporting the specific targeting of reconsolidation and the requirement of NA neuronal activity therein. The reconsolidation process begins shortly after memory recall and is complete within 6 hours [36, 37]. In the present study, CNO was administered immediately after memory recall and again 2 and 4 hours later to maintain neuronal inhibition throughout the duration of the reconsolidation process. Disruption of reconsolidation leads to a decrease in the retention of that specific memory and has been studied as a potential therapeutic strategy to ameliorate trauma-associated memories in patients with post-traumatic stress disorder [38, 39].

Using fosTRAP2XAi14 mice that had an established cocaine place preference, neurons that were activated during the recall of cocaine contextual memory were ‘trapped’ for further study. Results demonstrate that NA MSN in both the core and shell regions were activated by recall of cocaine contextual memories. We furthered the analysis of MSN activated by cocaine memory recall by examining the presence of neuroplastic changes. Analysis of MSN ‘trapped’ following cocaine memory reactivation showed changes indicative of synaptic strengthening (increased total spine density) and eventual maturation of synapses (increase in density of thin spines). Thin spines are highly dynamic (compared to mushrooms) and believed to represent “learning spines”, responsible for forming new memories [40]. The presence of these changes was also evident when examining measures of spine surface area, neck diameter and head diameter. Greater surface area following reactivation of cocaine memory indicates the rapid expansion of spines that occured following recall vs no memory reactivation. Reductions in spine neck and head diameter lend further support that the increase in spine density was driven by thin spines, whose characteristics include smaller head and neck diameter compared to other spine types. Prior literature on dendritic spines and cocaine in rodent models describes cocaine-induced changes in thin spines. For example, an increase in the number of the thin spine subtype, without changes in other spine subtypes, was found in the medial prefrontal cortex of rats following 7 days of forced abstinence from cocaine vs saline iv self-administration [41]. Interestingly, in a study assessing dendritic spine density of D1R- and D2R-expressing MSNs, increased spine density was observed in both MSN phenotypes when measured 2 days after chronic cocaine administration, while the increased spine density was maintained only in D1R-expressing MSNs 30 days after the last cocaine injection [41, 42]. In contrast to prior reports [41–43], our study measured dendritic plasticity induced by reactivation of cocaine memories and not by cocaine itself. Both experimental groups of mice had identical cocaine exposure and differed only by cocaine memory reactivation achieved by re-exposure to the cocaine-paired context or no memory reactivation. Thus reactivation of cocaine memories induced changes in spine morphology. In the current study, spine density was not evaluated in phenotypically defined MSN, so it remains unknown if recall of cocaine memories produced dendritic changes in D1R- and/or D2R-expressing MSN.

Results presented here demonstrated that cocaine memory reconsolidation is dependent on NA neuronal activation. Activation of NA neurons is accompanied by MSN dendritic plasticity suggesting increases in efferent connections to these neurons. To identify other components of the cocaine memory engram that may synapse on NA neurons, the role of NA projection neurons from the vHPC was investigated. This is the first study to demonstrate the vHPC to NA projection as necessary for the reconsolidation of cocaine contextual memories. The vHPC sends excitatory glutamatergic projections to the NA [44]. This excitatory projection has been implicated in encoding contextual information of addictive behaviors [45]. Furthermore, cue-induced recall of cocaine memory depends on particular projections from the vHPC to the NA (i.e., ventral CA1→NAcore) [17]. The vHPC is also responsible for mediating associative processing of cues during classical fear conditioning [46]. Finally, inhibition of the vHPC (with GABA receptor agonists) attenuates reinstatement to cocaine seeking following a presentation of either drug-paired cues or a priming injection of cocaine in a rat cocaine self-administration procedure [47].

Prior literature described the engram that mediates consolidation of cocaine CPP as being comprised of the ventral CA1 region of the hippocampus and the NAcore in mice [48]. In that study, chemogenetic silencing of either NAcore or vCA1 engram cells prior to the test for cocaine place preference impaired the expression of a cocaine CPP, suggesting these cells are involved in either the storage and/or retrieval of the CPP memory [48]. Their study differed from ours in several ways. Most importantly for our focus on reconsolidation, we administered CNO after, not before, memory reactivation and CPP expression was tested 3 days later without further CNO administration. Chemogenetic inhibition of vHPC neurons projecting specifically to the NA after cocaine memory recall interfered with reconsolidation as evidenced by erasure of the established cocaine place preference. Recall of cocaine contextual memories resulted in neuroplastic changes in the dendrites of activated pyramidal neurons in the ventral hippocampus. Changes in the density and morphology of dendritic spines reflect alterations in synaptic strength, particularly of excitatory synapses. Increases in dendritic spine density was noted and, in contrast to the nucleus accumbens where thin spines were most affected, in vHPC neurons the densities of all spine types were higher after cocaine memory recall.

Our prior studies using a similar mouse CPP model of cocaine contextual memory processes identified a cellular signaling pathway that is critical for the reconsolidation of cocaine memories. Recall of cocaine contextual memories resulted in activation of glycogen synthase kinase 3β (GSK3β) and its downstream effector, mTORC1, and inhibition of either was sufficient to block cocaine memory reconsolidation [21, 23]. Further investigation demonstrated that NMDA receptor activation, specifically GluN2A/B subtypes, is upstream of the activation of GSK3β in the NA and hippocampus, and activation of these NMDA receptors is a requirement for cocaine memory reconsolidation to occur [22]. Thus, it is possible to hypothesize based on the current findings and our previous work that cocaine memory reconsolidation involves activation of vHPC glutamatergic neurons projecting to the NA which stimulates NMDA receptors located on MSN and signaling through GSK3β and mTORC1 to produce changes in protein synthesis needed for long-term memory [49]. The literature supports glutamatergic synapses as key cellular sites where cocaine experience creates memory traces that can then promote cocaine craving and seeking, and that these synapses can be generated in the NA following cocaine experience [50].

What was not addressed in the current investigation is the identification of the target brain region(s) of the NA MSN involved in cocaine memory reconsolidation. The MSNs of the NA project to several brain areas including the ventral pallidum, the mesencephalon including the ventral tegmental area, and the hypothalamus [51]. In other work, it was demonstrated that D1 receptor-expressing MSNs largely collateralize to both the ventral pallidum and the ventral mesencephalon. It was further demonstrated that activation of the MSNs specifically innervating the ventral pallidum was necessary for cue-induced cocaine reinstatement in rats [52]. Thus, the ventral pallidum may be a target of the MSNs activated by and necessary for reconsolidation of cocaine memories. Further studies are needed to confirm this proposed circuit.

## Conclusions

Cue-induced relapse is a major barrier for successful remission in patients battling cocaine addiction and there are no FDA approved medications to treat cocaine use disorder [53–56]. Targeting the reconsolidation of cocaine contextual memories may be an important strategy to help prevent relapse and reduce drug seeking behaviors. Future directions include identifying the output pathway from the NA involved in the reconsolidation of cocaine contextual memories, as well characterizing possible sex differences in the circuitry responsible for the reconsolidation of cocaine contextual memories.

## Acknowledgments

We thank Joseph Meissler for management of the mouse breeding colony that generated the TRAP2XAi14 mice for this study, Chongguang Chen for his assistance in setting up imaging protocols, and Thomas Campion for viral vector preparation.

